# γδ intraepithelial lymphocytes facilitate pathological epithelial cell shedding via CD103-mediated granzyme release

**DOI:** 10.1101/2021.01.20.427150

**Authors:** Madeleine D. Hu, Natasha B. Golovchenko, Thomas J. Kelly, Jonathan Agos, Matthew R. Zeglinski, Edward M. Bonder, Inga Sandrock, Immo Prinz, David J. Granville, Alastair J.M. Watson, Karen L. Edelblum

**Affiliations:** Center for Immunity and Inflammation, Department of Pathology, Immunology & Laboratory Medicine, Rutgers New Jersey Medical School, Newark, NJ, 07103, USA; Department of Pathology and Laboratory Medicine, University of British Columbia, Vancouver, British Columbia, V6T 2B5, Canada; Department of Biological Sciences, Rutgers University – The State University of New Jersey, Newark, NJ, 07102, USA; Institute of Immunology, Hannover Medical School, Hannover, Germany; Institute of Systems Immunology, University Medical Center Hamburg-Eppendorf, 20246 Hamburg, Germany; Department of Gastroenterology and Gut Biology, Norwich Medical School, University of East Anglia, Norwich, UK

## Abstract

Excessive shedding of enterocytes into the intestinal lumen is observed in inflammatory bowel disease and is correlated with disease relapse. However, the mechanisms underlying this phenomenon remain unclear. Intraepithelial lymphocytes (IEL) expressing the γδ T-cell receptor (TCR) provide surveillance of the intestinal mucosa at steady-state, which is regulated, in part, by CD103. Intravital microscopy of lipopolysaccharide (LPS)-treated mice revealed that γδ IELs make extended contact with shedding enterocytes. These prolonged interactions require CD103 engagement by E-cadherin, as CD103 blockade significantly reduces LPS-induced shedding. Furthermore, we find that granzymes A and B, but not perforin, are required for cell shedding, and that these granzymes are released by γδ IELs both constitutively and following CD103/E-cadherin ligation. These findings indicate that extracellular granzyme facilitates shedding, likely through cleavage of extracellular matrix proteins. Our results uncover a previously unrecognized role for γδ IELs in facilitating pathological cell shedding in a CD103- and granzyme-dependent manner.

## Introduction

The intestinal epithelium is a single cell layer that serves as a critical barrier to prevent interaction between luminal contents and the mucosal immune system^1,2^. The maintenance of the small intestinal epithelial barrier is dependent upon tight regulation between proliferation in the crypt and shedding, or extrusion of epithelial cells, at the villus tip^3^. While physiological cell shedding at steady-state is required to maintain epithelial homeostasis, the rate of shedding is significantly enhanced in inflammatory bowel disease (IBD)^4-7^. In this context, pathological shedding results in the loss of multiple contiguous epithelial cells, which can lead to ulceration or allow entry of luminal antigen and bacteria into the underlying tissue, thus inducing inflammation.

IBD is a multi-factorial disease that affects 3.1 million Americans. Genetic factors, barrier defects, immune dysregulation and microbial dysbiosis are all thought to contribute to disease susceptibility^8^. Overproduction of tumor necrosis factor (TNF) is a hallmark of intestinal inflammation in IBD^9,10^, and therefore, biologic therapy using anti-TNF antibodies is widely used in the treatment of these patients^11-13^. While approximately one-third of patients respond well to anti-TNF therapy and achieve clinical remission, one-third of patients exhibit only an intermediate response, and the remaining one-third are non-responsive^14^. Since a substantial increase in epithelial cell shedding is a known predictor of relapse in Crohn’s disease patients^15-17^, it may be possible to enhance or supplement the clinical response to anti-TNF therapy in intermediate responders by identifying additional mechanisms that regulate epithelial cell shedding.

Pathological cell shedding can be modeled in mice through the systemic administration of pro-inflammatory mediators such as TNF or lipopolysaccharide (LPS)^18,19^. Both of these stimuli have been shown to induce excessive cell shedding in mice through an epithelial TNFR1-dependent signaling mechanism. These TNF-mediated shedding events can be identified by caspase-3 activation and the formation of an actomyosin funnel that surrounds the shedding cell as it is extruded^19^. Nuclear fragmentation is observed as the cell is extruded into the lumen. While cytoskeletal remodeling and caspase activation are critical for the shedding process^19,20^, much remains unknown regarding the mechanisms by which individual cells are targeted for extrusion or the contributions of direct immune cell contacts to this process.

Intraepithelial lymphocytes (IELs) expressing the γδ T cell receptor (TCR) bridge innate and adaptive immunity, exhibit a largely protective response to dampen acute inflammation^21,22^, and promote mucosal barrier integrity^23-25^.

Further, γδ IELs also contribute to epithelial turnover and proliferation in response to injury^21,22,26^. Although proliferation and shedding are inextricably linked, the role of γδ IELs in cell shedding has yet to be investigated. Despite epithelial cells outnumbering IELs by 10:1, we have shown that γδ IELs provide immune surveillance of the villous epithelium by engaging in dynamic patrolling behaviors^27,28^. γδ IELs migrate along the basement membrane (BM) and between adjacent epithelial cells in the lateral intercellular space (LIS).^27^ This motility is dependent upon ligand-binding interactions between the IEL and enterocyte, with homotypic occludin interactions and CD103 (α_E_β_7_ integrin)/E-cadherin binding being critical for the motility and retention of γδ IELs within the LIS, respectively.

Although epithelial apoptosis can be induced through cytolytic functions of IELs, which are largely CD8^+^, the role of γδ IELs in pathological cell shedding has yet to be investigated. Herein, we have used intravital microscopy to demonstrate that γδ IELs make prolonged contacts with shedding epithelial cells prior to their extrusion in LPS-treated mice. These extended cell-cell contacts are dependent upon CD103 ligation, and both genetic ablation and antibody-mediated blockade of CD103 reduces the severity of LPS-induced cell shedding. Further, we show that granzymes A and B, which are released by γδ IELs both at steady-state and in response to CD103 engagement, are required for pathological cell shedding. We find that this process is perforin-independent, suggesting that CD103 ligation stimulates the extracellular release of granzymes by γδ IELs that contribute to cell shedding likely through the proteolytic cleavage of extracellular matrix proteins. These findings represent a previously undiscovered role for γδ IELs in facilitating cell shedding and may provide a novel therapeutic target for maintenance of mucosal homeostasis in IBD.

## Results

### γδ IELs directly interact with shedding cells in response to pathological concentrations of lipopolysaccharide

Previously published electron microscopy studies observed IELs located directly beneath or adjacent to shedding enterocytes under homeostatic conditions^29,30^. While it was hypothesized that IELs may plug the gap left behind by the extruded cell, recent studies provide conclusive evidence that neighboring epithelial cells extend protrusions underneath the shedding cell to maintain an intact epithelial monolayer and prevent an influx of luminal antigen into the underlying mucosal compartment^19,31^. Although these studies have elucidated many of the epithelial cell-intrinsic intracellular signals involved in coordination of the shedding process, whether IELs are actively involved in the shedding of enterocytes into the lumen remains unclear.

To determine whether IELs contact shedding cells in the context of inflammation, we treated wildtype (WT) mice with LPS for 90 min to rapidly induce epithelial cell shedding, as quantified by cellular morphology and cleaved caspase-3 (CC3)-positive staining (Fig. 1A,B)^18^. We then used transmission electron microscopy to examine jejunal villi from LPS-treated mice (Fig. 1C) and identified multiple instances in which IELs were in direct contact with shedding events. To investigate the frequency of IEL contact with shedding cells, we treated mice expressing a Tcrd-histone 2B enhanced GFP reporter (TcrdEGFP)^32^ with LPS and measured the proximity of GFP^+^ γδ IELs to CC3^+^ shedding events (Fig 1D). Overall, shedding epithelial cells were more likely to have a γδ IEL within a two epithelial cell distance compared to a CC3^-^ enterocyte (p<0.05, Fig. 1E).

**Figure 1.**
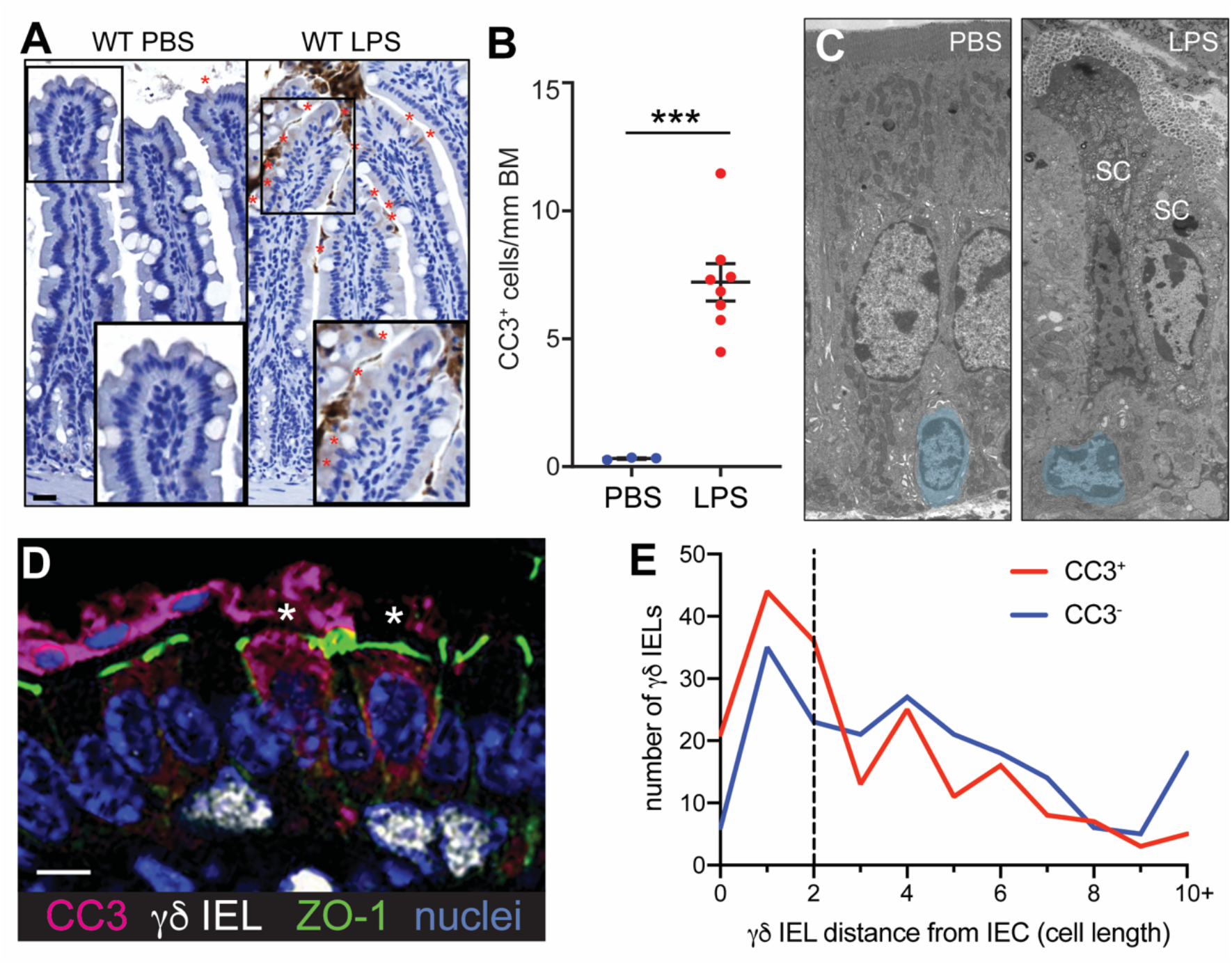
γδ IELs directly interact with shedding enterocytes. (A) Immunohistochemical staining and (B) quantification of cleaved caspase-3 (CC3)-positive shedding events in the jejuna of PBS- or LPS-treated WT mice (marked by red asterisks). Scale bar = 50 μm. (C) Representative transmission electron micrographs of PBS- and LPS-treated WT jejuna. Intraepithelial lymphocytes are pseudocolored blue. SC, shedding cell. (D) Immunofluorescent micrograph of γδ IELs (white) interacting with CC3^+^ shedding cells (magenta). Green:ZO-1; blue: nuclei. Asterisks indicate shedding cells. Scale bar = 5 μm. (E) Quantification of distance (number of epithelial nuclei) between a CC3^+^ shedding or CC3^-^ enterocyte and the nearest γδ IEL. n = 3-8 mice per condition; ***p<0.001.

We next performed intravital microscopy on the jejunal mucosa of LPS-treated TcrdEGFP mice to examine the spatiotemporal dynamics of IEL-shedding enterocyte interactions. This technique enabled us to observe shedding cells as they were extruded into the lumen as well as the characteristic surveillance behavior of γδ IELs^27^. We found that approximately 46% of shedding cells were contacted by a γδ IEL prior to their extrusion (Fig. 2A, Video S1), and that the γδ IELs that made contact with shedding events migrated at slower speeds relative to γδ IELs that did not (Fig. 2B).

**Figure 2.**
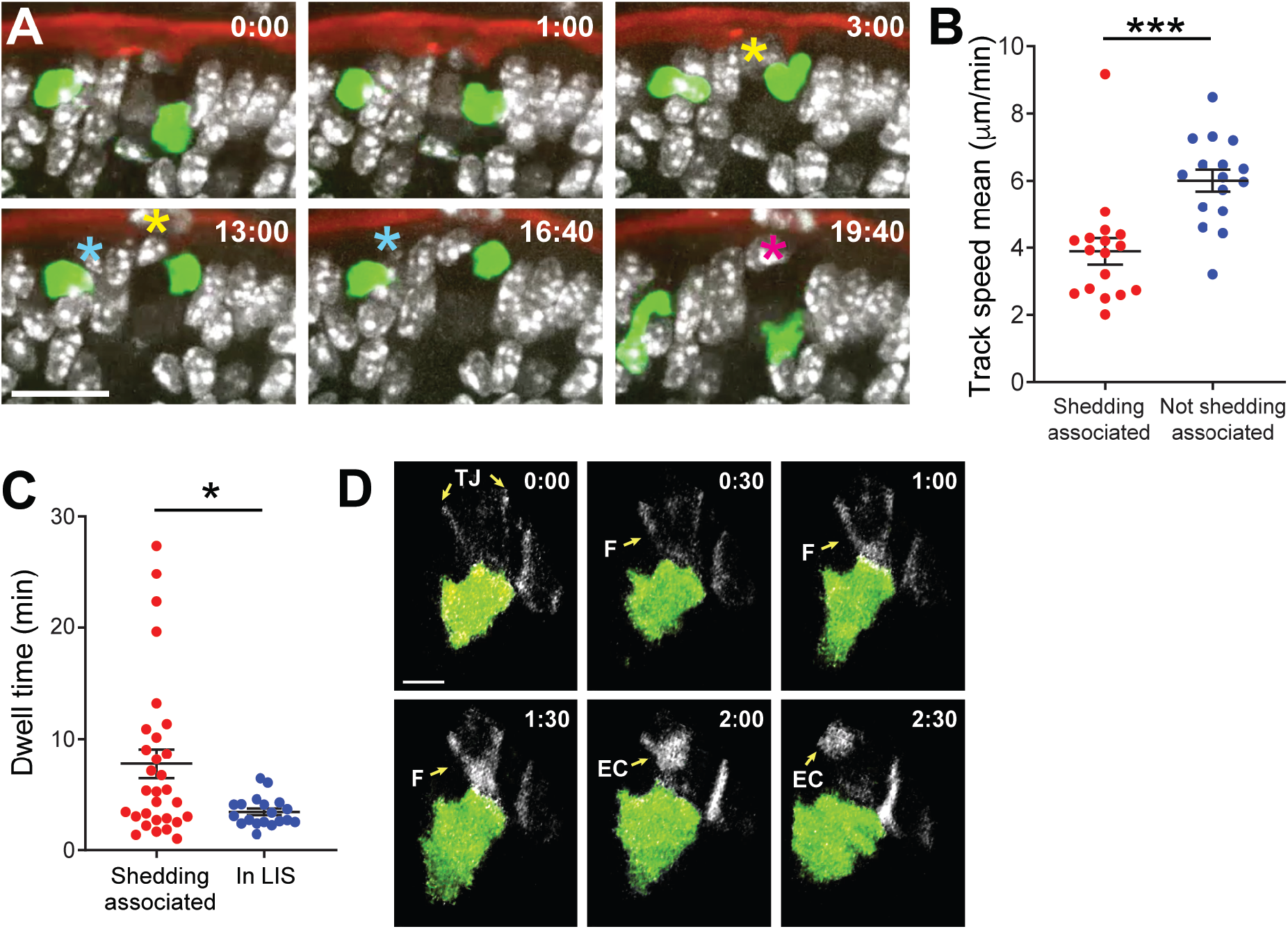
γδ IELs maintain prolonged contact with enterocytes prior to extrusion into the lumen. Intravital microscopy of (A) TcrdEGFP or (D) TcrdGDL mRFP-ZO1 mice treated with LPS. (A) Time lapse images from intravital imaging show γδ IELs (green) interacting with multiple shedding cells (asterisks). Red: lumen; white: nuclei. Scale bar = 20 μm. (B) Mean track speed of γδ IELs that are or are not associated with shedding cells. (C) Dwell time of γδ IELs at sites of enterocyte shedding or located within the LIS. (D) γδ IEL (green) interacting with a ZO-1 funnel (F, white). TJ: tight junction; EC: extruding cell. Scale bar = 5 μm. n = 6-8 mice; *p<0.05, ***p<0.001.

This reduced migratory speed is likely attributable to the increased dwell time of γδ IELs at the base of shedding enterocytes, which was 56% longer than the average dwell time of γδ IELs in the LIS (7.8 vs. 3.4 min, respectively) (Fig. 2C). Interestingly, we did not observe CD8^+^ TCRαβ ^+^(GFP^-^) IELs associating with shedding events in LPS-treated mice, indicating that this change in migratory behavior may be unique to γδ IELs.

Based on these findings, we next asked whether γδ IELs also interact with shedding events under steady-state conditions. To address this, we analyzed 8 videos acquired from intravital imaging of untreated TcrdEGFP mice and quantified the frequency of γδ IEL interactions with shedding enterocytes in the jejunum. As expected, the relative frequency of cell extrusion in untreated mice was rare; however, we found that nearly a third of these shedding events were contacted by a γδ IEL immediately prior to expulsion into the lumen.

Epithelial cell shedding in response to inflammatory stimuli is a well-defined process that begins with junction protein rearrangement and formation of an actin funnel and concludes with extrusion of the cell into the intestinal lumen^19^. To determine the point in this process at which γδ IELs contact the shedding cell, intravital microscopy was performed on mice expressing both a Tcrd cytoplasmic eGFP reporter (TcrdGDL)^33^ and a monomeric RFP (mRFP)-zonula occludens 1 (ZO1) fusion protein^34^. These studies revealed that γδ IELs contact shedding cells prior to the formation of the actin funnel (Fig. 2D, Video S2), indicating that these IELs are present during or shortly after initiation of the shedding process. Taken together, these data demonstrate that γδ IELs maintain prolonged contact with epithelial cells beginning in the early stages of pathological cell shedding in response to LPS.

### CD103 expression is required for pathological cell shedding

The extended contact that we observed between γδ IELs and shedding enterocytes next led us to investigate whether the IEL dwell time at sites of cell shedding was critical to the extrusion process. We have previously shown that loss of CD103 increases γδ IEL motility by reducing dwell time in the LIS, likely through disruption of CD103/E-cadherin interactions^27,28^. Thus, to determine whether altering the duration of IEL/epithelial contact affects pathological cell shedding, we investigated the extent of cell extrusion in LPS-treated WT and CD103-deficient (CD103 KO) mice. At the peak of shedding, CD103 KO mice exhibited considerably reduced rates of shedding relative to WT mice in all three regions of the small intestine, with a 69% reduction observed in the jejunum. (Fig. 1A, 3A,B, S1A). Loss of CD103 also abrogated LPS-induced shedding at later timepoints, indicating that the marked reduction in CC3^+^ cells was not due to a delayed response (data not shown). Similarly, CD103 KO mice exhibited a 68% reduction in shedding relative to WT in response to pathological concentrations of TNF (Fig. 3C), demonstrating that loss of CD103 confers protection against TNFR-dependent shedding events^18,19^. Notably, villi in CD103 KO mice were longer than those in WT mice, further supporting a role for γδ IELs in physiological cell shedding (Fig. S1B). We next asked whether the loss of CD103 resulted in stalling of the shedding process by analyzing the number of ZO-1 funnels and/or CC3^+^ enterocytes in LPS-treated and WT and CD103 KO mice. If CD103 was required for caspase activation, we would expect to observe an increase in the number of incomplete shedding events since actomyosin contraction can occur in the absence of caspase activation^19^; however, this is not the case (CC3^-^ funnels, Fig S1C). Together, these data support our observation that γδ IELs directly contact enterocytes during the initial stage of shedding, an interaction dependent upon CD103 ligation with epithelial E-cadherin.

**Figure 3.**
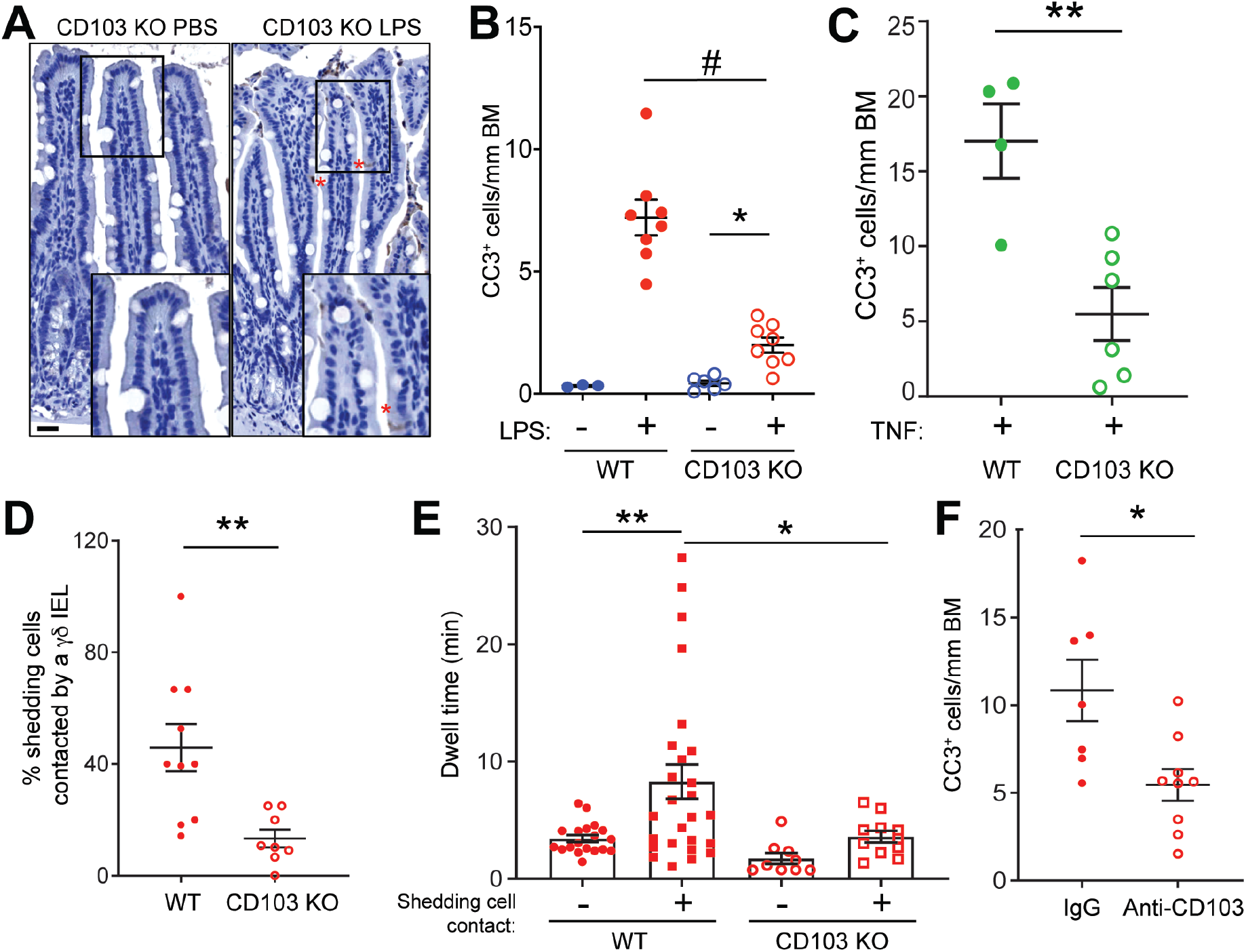
Deletion or antibody-mediated blockade of CD103 abrogates pathological cell shedding. Representative CC3 immunostaining and quantification of (A,B) CC3^+^ epithelial cells in WT or CD103 KO mice treated with LPS or (C) TNF. Asterisks: shedding cells. Scale bar = 50 μm. (D) Percent of shedding cells contacted by a γδ IEL and (E) γδ IEL dwell time near shedding cells in LPS-treated WT or CD103 KO mice. (F) Quantification of CC3^+^ shedding cells in the jejunum of WT mice treated with anti-CD103 or isotype control prior to LPS administration. n = 3-9 mice (9-26 tracks); *p<0.05, **p<0.01, #p<0.0001.

To determine whether differences in the microbiome contributed to the reduced cell shedding observed in CD103 KO mice, WT and CD103 KO mice were cohoused after weaning and then administered LPS at 8 weeks of age. Cohousing had no appreciable effect on the protection against cell shedding conferred by loss of CD103 expression (Fig. S1D). Although CD103 promotes migration of γδ IELs and other TNF-producing immune cells into and within the gut, ELISA performed on whole intestinal lysates from LPS-treated WT and CD103 KO mice showed that TNF production was similar between the two (data not shown). These findings indicate that the decrease in LPS-induced cell shedding observed in CD103-deficient mice cannot be attributed to reduced mucosal TNF production or an altered microbiome.

To evaluate γδ IEL interactions with the few shedding events that do occur in absence of CD103, we performed intravital microscopy on LPS-treated TcrdEGFP; CD103 KO mice. Time-lapse video microscopy showed that γδ IELs contact shedding events in CD103 KO mice less frequently than in WT (Fig. 3D). Consistent with the role of CD103 in facilitating γδ IEL retention within the LIS, the few γδ IELs that contacted shedding enterocytes in CD103 KO mice migrated more rapidly and exhibited reduced dwell times near these shedding events as compared to their LPS-treated WT counterparts (Fig. 3E, S1E).

Next, we investigated whether inhibition of CD103 interaction with its binding partner, E-cadherin, is sufficient to reduce cell shedding. To test this, an anti-CD103 blocking antibody (M290) was administered prior to treatment with LPS (Fig. S1F). Mice treated with anti-CD103 exhibited a 50% decrease in LPS-induced cell shedding compared to mice receiving an isotype control (Fig. 3F). Similar to our findings in CD103 KO mice, these reductions in LPS-induced cell shedding were not accompanied by reduced TNF production (data not shown). Together, these data demonstrate a requirement for CD103 binding in the cell shedding process and show that short-term blockade of CD103 is sufficient to reduce the frequency of shedding events.

To determine whether CD103 expressed specifically on γδ IELs was responsible for facilitating pathological cell shedding, we generated mixed bone marrow chimeras by engrafting lethally-irradiated γδ T-cell-deficient (Tcrd KO) mice with 80% Tcrd KO and 20% CD103 KO or WT bone marrow. While we were able to reconstitute the γδ IEL compartment in mice receiving WT bone marrow, CD103-deficient γδ T cells were unable to repopulate the IEL niche to a comparable level. As a result, we were unable to directly assess whether the effect of CD103 on cell shedding was directly attributable to γδ IELs. However, our findings demonstrate a novel role for CD103 in mediating pathological cell shedding and show that antibody-mediated blockade of α_E_ integrin can reduce the severity of the shedding response.

### γδ IELs are not required for LPS-induced cell shedding

Previous studies have demonstrated that TNF induces cell shedding through a mechanism independent of adaptive immunity^35^. These experiments were performed in *Rag*- and *Tcrd-Tcrb*-deficient animals; however, the specific contribution of γδ T cells has not been addressed. To this end, we first quantified LPS-induced cell shedding in WT and Tcrd KO mice and observed no difference in the frequency of cell shedding (Fig. 4A). Since the loss of γδ T cells can be partially compensated by the expansion of other immune cells with similar functions^33^, we next evaluated LPS-induced cell shedding following conditional depletion of γδ T cells. Mice expressing the diphtheria toxin receptor (DTR) downstream of the Tcrd promoter (TcrdGDL)^33^ were treated with DT to deplete γδ T cells without affecting baseline cell shedding rates (Fig. S2). The number of LPS-induced shedding events was similar between DT- and vehicle-treated TcrdGDL mice, as well as DT-treated WT controls (Fig. 4B). Overall, these results suggest that despite the direct contact we have observed between γδ IELs and shedding cells, γδ T cells are not required for LPS-induced cell shedding.

**Figure 4.**
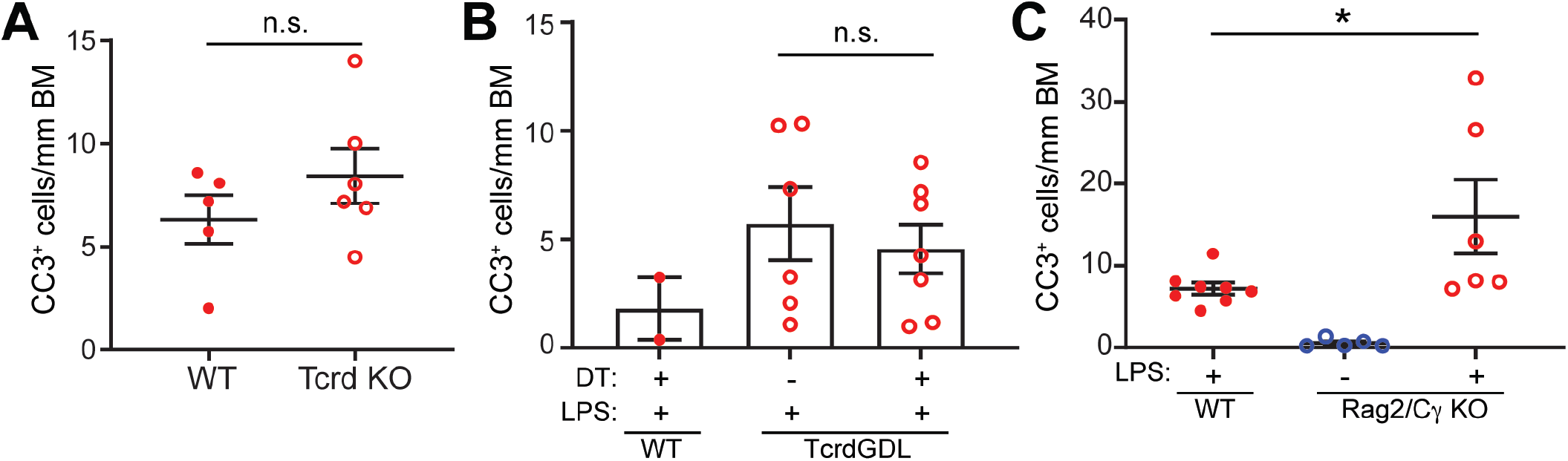
γδ IELs are not required for LPS-induced cell shedding. Quantification of CC3+ shedding events in jejunal tissue of (A) WT or Tcrd KO mice treated with LPS. (B) WT or TcrdGDL mice were treated with diphtheria toxin (DT) or vehicle control prior to LPS treatment. (C) WT or Rag2/Cγ KO mice were treated with PBS or LPS. n = 2-8 mice; *p<0.05, n.s., not significant.

CD103 is expressed by a variety of different immune cells in the intestinal mucosa, including IELs and intraepithelial innate lymphoid cells (ILC). We therefore asked whether ILCs are required for pathological cell shedding by treating Rag2/Cγ KO mice, which are deficient for B cells, T cells, and ILCs, with LPS. The extent of cell shedding was similar as to that observed in WT mice (Fig. 4C), demonstrating that pathological epithelial cell shedding can occur independently of both adaptive immune cells and ILCs. While these lymphocytes are not required for cell shedding to occur, our findings clearly indicate that both CD103 expression and the interactions between γδ IEL and shedding cells can contribute to this process.

### The recognition of self- or stress ligands do not contribute to γδ IEL-mediated LPS-induced cell shedding

We hypothesized that γδ IEL recognition of shedding cells via ligand-receptor interactions is involved in promoting the extended contact between IELs and shedding epithelial cells. Multiple studies have suggested that the γδ T cell receptor (TCRγδ) can recognize self-antigens, some of which may be upregulated by cell stress ^36-38^, thus we first investigated the requirement for TCRγδ in LPS-induced cell shedding by inducing TCR internalization in WT mice with an anti-TCRγδ antibody^39^. However, anti-TCRγδ treatment had no effect on cell shedding in response to LPS (Fig. S3A), indicating that signaling through the TCRγδ is dispensable for this process. These findings are consistent with a previous study demonstrating that TCRγδ signaling is not required for γδ IEL migratory behavior in the context of infection^40^. Next, we investigated a potential role for NKG2D-mediated innate-like recognition of stress ligands in pathological cell shedding. We found that LPS treatment did not induce the expression of epithelial stress ligands recognized by NKG2D (Fig. S3B-D). Furthermore, pre-treatment of mice with an anti-NKG2D blocking antibody failed to abrogate pathological cell shedding (Fig. S3E). Thus, the mechanism by which γδ IELs detect shedding cells remains an ongoing area of investigation.

### Granzymes mediate LPS-induced epithelial cell shedding

In addition to enabling ligand-binding interactions between IELs and epithelial cells, we posited that prolonged retention of IELs at sites of cell extrusion may also allow these cells to secrete a high local concentration of soluble proteins to facilitate shedding. Granzymes (Gzm) are a family of proteases with a wide range of targets that are highly expressed by NK cells, cytotoxic lymphocytes, and γδ IELs^41,42^. In conjunction with perforin, a pore-forming protein that facilitates granzyme entry into target cells, granzymes initiate intracellular signaling cascades that can ultimately result in apoptosis^43^. Moreover, signaling through CD103 has been shown to promote degranulation of cytotoxic lymphocytes, resulting in the local polarized release of granzymes^44,45^. Therefore, to investigate the requirement for granzyme expression in LPS-induced cell shedding, lethally-irradiated Tcrd KO mice were engrafted with WT or GzmA/B-deficient bone marrow, thus generating bone marrow chimeras in which all γδ T cells are GzmA- and B-deficient. We found that the initial irradiation of these mice did not affect baseline cell shedding rates (data not shown), and the resultant chimeras exhibited similar proportions of γδ and αβ IELs, regardless of the donor genotype (Fig. S4). Following LPS treatment, mice engrafted with GzmA/B-deficient bone marrow exhibited a 31% decrease in cell shedding rates compared to WT, suggesting a role for at least one of these granzymes in mediating LPS-induced cell shedding (Fig. 5A).

**Figure 5.**
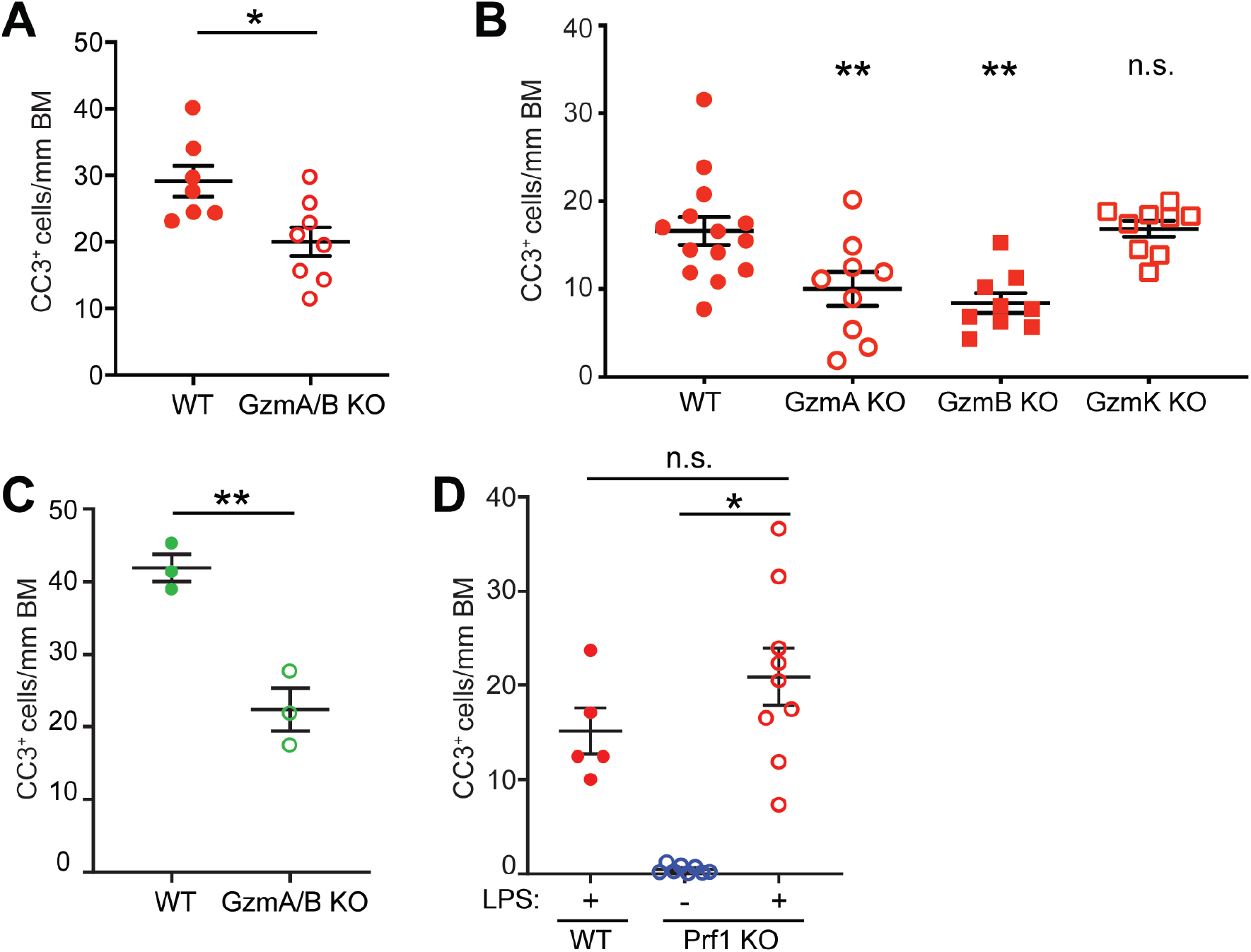
Granzymes A and B mediate LPS-induced epithelial cell shedding in a perforin-independent manner. Quantification of CC3^+^ shedding events in jejunal tissue. Lethally irradiated Tcrd-deficient mice were engrafted with (A) GzmA/B DKO or (B) WT, GzmA KO, GzmB KO, GzmK KO and treated with LPS. (C) GzmA/B KO or WT mice were treated with TNF. (D) WT or Prf1 KO mice were treated with PBS or LPS. n = 3-14 mice; n.s., not significant *p<0.05, **p<0.01.

To examine the roles of individual granzymes in LPS-induced cell shedding, bone marrow chimeras were made with Tcrd KO mice engrafted with WT, GzmA-, GzmB-, or GzmK-deficient bone marrow. While GzmK-deficient bone marrow chimeras did not exhibit significant differences in shedding rates as compared to WT-engrafted mice, GzmA- or GzmB-deficient chimeras displayed 62% and 55% reductions in LPS-induced cell shedding, respectively (Fig. 5B), indicating that GzmA and B both contribute to pathological cell shedding. Both of these granzymes have been implicated in stimulating TNF production downstream of LPS-induced TLR4 activation^46,47^. To test whether this phenomenon was responsible for the reduction in LPS-induced cell shedding observed in GzmA/B KO bone marrow chimeras, we treated GzmA/B-deficient mice with high-dose TNF. We found that GzmA/B KO mice exhibit a 47% decrease in shedding in response to TNF (Fig. 5C), indicating that the effect that with LPS treatment was not solely due to the potential pro-inflammatory role of these enzymes.

Based on this requirement for GzmA and B in LPS-induced shedding, we next asked whether γδ IELs directly induce epithelial apoptosis through a perforin-dependent mechanism. To address this, we quantified cell shedding in LPS-treated perforin-deficient mice (Prf1 KO) and found that the frequency of shedding events in these mice were similar to those observed in WT mice (Fig. 5D). Overall, these data indicate that GzmA and B promote LPS-induced shedding through a pathway independent of perforin-mediated intracellular enzymatic activity.

### CD103 ligation mediates γδ IEL granzyme release

Having demonstrated that GzmA and B facilitate LPS-induced cell shedding and given that these enzymes are among the most highly expressed proteins in γδ IELs at steady-state^42,48^, we hypothesized that γδ IELs participate in pathological cell shedding via granzyme release into the extracellular space. Thus, to investigate the mechanisms by which γδ IELs secrete granzymes, we first assessed GzmA and B release by untreated, sort-purified WT γδ IELs. We were surprised to find that both GzmA and B are constitutively secreted by γδ IELs, with GzmA released to a greater extent than GzmB (375 pg vs. 1.5 pg/10^5^ cells). Since the interaction between CD103 and E-cadherin has been shown to promote degranulation of cytotoxic lymphocytes ^44,45^, we asked whether CD103/E-cadherin binding could promote granzyme secretion by stimulating freshly-isolated γδ IELs with E-cadherin-Fc (E-cad-Fc). Notably, E-cad-Fc stimulation induced granzyme secretion, albeit to a lesser extent than anti-CD3 (Fig. 6A,B). Whereas anti-CD3 treatment resulted in substantial granzyme release via degranulation, as indicated by externalization of LAMP1 (CD107a), very little LAMP1 was observed on the surface of E-cad-Fc-stimulated cells (Fig. 6C). Consistent with the degranulation-independent release of granzyme following E-cad-Fc engagement, immunofluorescence of freshly-isolated γδ IELs revealed a subset of GzmA- or GzmB-containing vesicles that did not co-localized with LAMP1 (Fig. 6D). Taken together, this data indicates that γδ IELs secrete GzmA and B constitutively at steady-state and following CD103 engagement, which further potentiates granzyme release through LAMP1^-^ secretory vesicles.

**Figure 6.**
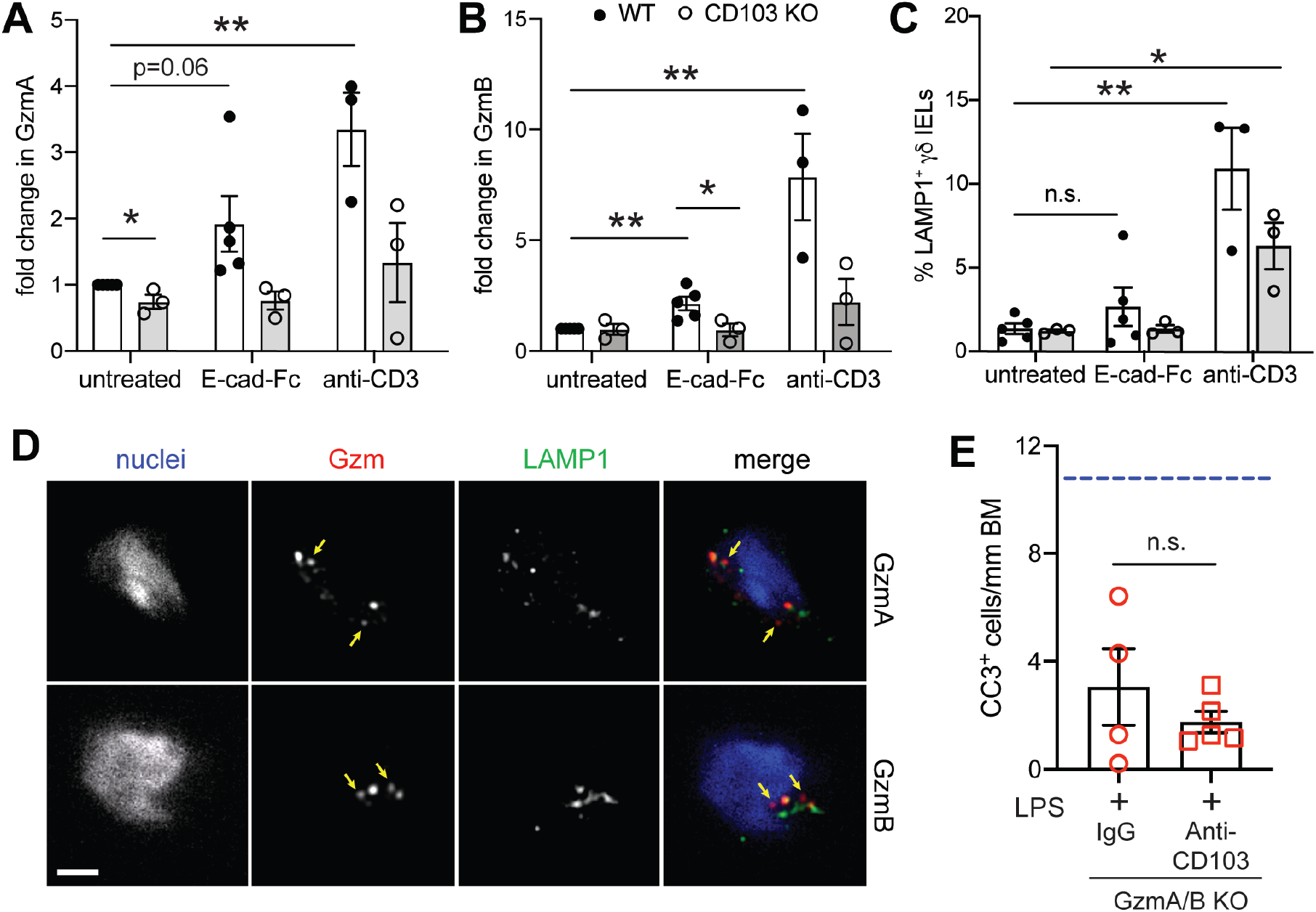
CD103 ligation promotes degranulation-independent granzyme release by γδ IELs. γδ IELs were isolated from WT or CD103 KO mice, sort-purified, stimulated with E-cad-Fc or anti-CD3 and ELISA performed on cell supernatants for (A) GzmA and (B) GzmB. Data are shown as the fold change compared to untreated WT γδ IELs. (C) Frequency of LAMP1 externalization in IELs under the same treatment conditions. (D) Immunofluorescence micrographs of WT γδ IELs showing LAMP1 (green), GzmA or B (magenta), and nuclei (white). Yellow arrows indicate GznT^+^ LAMPT^-^ vesicles.(E) Quantification of CC3^+^ shedding cells in LPS-treated GzmA/B KO mice pre-treated with IgG or anti-CD103. Dashed line indicates representative amount of shedding observed in LPS-treated WT mice. n.s., not significant *p<0.05, **p<0.01.

Based on these findings, we next asked whether CD103 and GzmA/B contribute to LPS-induced shedding through the same pathway. To this end, GzmA/B KO mice were treated with LPS and either anti-CD103 or IgG control antibody and shedding was quantified by CC3 staining. We find that blocking CD103 engagement does not further reduce the extent of pathological cell shedding in GzmA/B-deficient mice (Fig. 6E), thus supporting a model in which granzyme release occurs downstream of CD103/E-cadherin ligation to facilitate cell extrusion.

## Discussion

We now describe a novel role for γδ IELs in facilitating pathological epithelial cell shedding, a phenomenon that can lead to the disruption of epithelial barrier integrity under inflammatory conditions. Our findings show that γδ IELs exhibit prolonged contact with shedding enterocytes prior to their extrusion into the lumen and that genetic or antibody-mediated inhibition of CD103/E-cadherin interactions abrogate cell shedding in response to LPS. Further, loss of CD103 expression significantly reduces the frequency of TNF-induced shedding events. These findings are consistent with our previous reports demonstrating that CD103 binding modulates the duration of γδ IEL/epithelial interactions^27,28^ and suggests that prolonged contact between γδ IELs and enterocytes fated to be shed may be central to the extrusion process. In regard to the mechanism by which γδ IELs facilitate cell shedding, we find that γδ IELs constitutively secrete both GzmA and B, which is enhanced following CD103 engagement by E-cadherin. Mice lacking GzmA and/or GzmB, but not perforin, exhibit reduced LPS-induced shedding, thus suggesting a role for extracellular granzyme release in this process. Moreover, treatment with anti-CD103 failed to further inhibit LPS-induced shedding in the absence of GzmA and B, indicating that granzyme release likely occurs downstream of CD103 ligation. Taken together, our data suggest that CD103 ligation functions to prolong γδ IEL contact with enterocytes and promote granzyme secretion, allowing the IEL to facilitate epithelial cell shedding by local, granzyme-mediated remodeling of the underlying BM.

Given the cytolytic capacity of γδ IELs^49^, we investigated whether γδ IELs function as professional killers during the shedding process. Although IELs sense and kill infected or stressed enterocytes via activation of the TCR or recognition of stress ligands by NKG2D^37,38,50,51^, blocking either receptor had no effect on the frequency of shedding events. This was not surprising since changes in IEL surveillance behavior in response to bacterial invasion occur independently of TCR activation^52^. Furthermore, granzymes A and B contributed to LPS-induced cell shedding in a perforin-independent manner, providing strong evidence that γδ IELs facilitate cell shedding through a mechanism other than direct lysis. Extracellular granzyme has been shown to promote the proteolytic cleavage of ECM proteins such as laminin, type IV collagen, and fibronectin, which are found in the intestinal BM^53-58^. Extracellular GzmB was recently shown to cleave α_6_ integrin, a component of the hemidesmosome that is critical in anchoring epithelial cells to the BM^59^. Moreover, circulating lymphocytes have been shown to secrete GzmA and B in order to remodel the vascular BM and facilitate extravasation^54,55,60^. These reports lead us to hypothesize that γδ IELs promote cell shedding by remodeling the BM beneath cells fated to be shed. Given the role of GzmA and B in promoting lymphocyte migration^54,55,60^, future studies will investigate the possibility that granzymes contribute to cell shedding by facilitating γδ IEL motility.

Previous work from our laboratory has demonstrated that both ligand-receptor interactions as well as cytokine signaling can influence the dynamics of γδ IEL motility^27,28,61^. Although largely presumed to function as a gut homing and retention signal, the biological consequences of CD103 ligation by E-cadherin remain unclear. In the context of cell migration, CD103/E-cadherin interactions influence lymphocyte morphology and motility^62^, and deletion of CD103 reduces γδ IEL retention in the LIS, thus increasing the patrolling behavior of these cells to limit bacterial invasion^28^. CD103 also stabilizes the immunological synapse between CTLs and E-cadherin-expressing target cells, thus enhancing TCR-mediated CTL activation^63^. We now show that CD103 functions to retain γδ IELs at sites of cell shedding, which may facilitate the local accumulation of γδ IEL-secreted granzymes in the extracellular milieu, resulting in degradation of ECM proteins and remodeling of the BM to promote cell extrusion. Moreover, CD103 ligation by E-cadherin is sufficient to stimulate granzyme secretion in the absence of additional TCR signaling. This finding is consistent with a previous study showing that engagement of CD103 with E-cad-Fc induces granule polarization and granzyme release by CTLs^44^. However, the absence of LAMP1 externalization upon E-cad-Fc-stimulated granzyme release by γδ IELs reveals an alternative pathway for granzyme secretion, one that is reminiscent of the constitutive release of newly-synthesized granzymes by activated CTLs^64^. Together, these data reveal a dual role for CD103 in (1) mediating extended contact between γδ IELs and shedding enterocytes, and/or (2) directly stimulating granzyme release through engagement of epithelial E-cadherin, ultimately resulting in the enhanced local deposition of granzymes to facilitate cell shedding.

Despite our intravital imaging studies demonstrating clear interactions between γδ IELs and shedding enterocytes, we were unable to demonstrate a requirement for γδ IELs in the cell shedding process, as neither germline nor DT-induced deletion of γδ T cells affected the frequency of LPS-induced shedding events. This led us to consider the possibility that another CD103^+^, GzmA/B-expressing leukocyte was involved. However, we found that Rag2/Cγ-deficient mice, which lack T, B, and innate lymphoid cells ^65,66^ exhibited moderately elevated levels of LPS-induced cell shedding. This finding is consistent with a study by Lau et al. showing that Rag1- and Tcrd/Tcrb-knockout mice are both more susceptible to TNF-induced cell shedding^35^ due to reduced monocyte chemoattractant protein-1 (MCP-1) production and a concomitant influx of plasmacytoid dendritic cells (pDCs) into the lamina propria. Thus, a similar mechanism may drive the increased susceptibility to LPS-induced shedding observed in Rag2/Cγ-deficient mice and also explain why loss of γδ T cells does not affect the extent of cell shedding.

The association between pathological cell shedding and IBD relapse^17^ coupled with our findings regarding the involvement of CD103 in this process highlight a potential benefit for CD103 blockade in IBD therapy. Vedolizumab has been shown to induce and sustain disease remission by blocking α_4_β_7_ integrin, thus inhibiting the entry of lymphocytes into the gut^67-69^. However, the effect of this drug on the migratory behavior of tissue-resident lymphocytes such as IELs remains unknown. Etrolizumab, which targets the β_7_ subunit of both α_4_β_7_ and α_E_β_7_ (CD103) integrins, is currently in phase III clinical trials^70^. Given our studies implicating CD103 in pathological cell shedding, it stands to reason that a pan-β_7_ inhibitor may provide the additional benefits of regulating IEL motility and limiting shedding events. This hypothesis is supported by a previous study showing that blockade of mucosal addressin cell adhesion molecule-1 (MAdCAM-1), which inhibits recruitment of T cells into the mucosa following TNF treatment, does not alter TNF-induced shedding rates, suggesting that tissue-resident lymphocytes rather than circulating T cells participate in regulation of cell shedding^35^. Therefore, targeting the β_7_ integrin subunit may improve disease outcomes and maintenance of remission in IBD patients.

Taken together, our data show a novel role for γδ IELs in facilitating pathological epithelial cell shedding through extended, CD103-mediated cell-cell contacts and extracellular granzyme release. γδ IEL-derived granzymes may accumulate near cells that are fated to be shed and promote their detachment from the basement membrane via proteolytic remodeling of the extracellular matrix. Our findings, particularly those indicating that antibody-mediated CD103 blockade can limit pathological cell shedding, strengthen the therapeutic potential of CD103 and suggest that drugs such as etrolizumab could be applied towards controlling cell shedding via modulation of γδ IEL behavior in IBD patients in remission, with an ultimate goal of maintaining homeostasis and preventing relapse.

## Materials and methods

### Animals

Mice of both sexes were used at 8-13 weeks of age and maintained on a C57BL/6 background under specific pathogen-free (SPF) conditions. Wildtype, Tcrd knockout (KO)^71^, and Prf1 KO mice were obtained from the Jackson Laboratory. TcrdH2BEGFP (TcrdEGFP) mice^32^ were provided by Bernard Malissen (INSERM) and crossed to CD103 KO (Jackson Labs) or mRFP-ZO1 Tg mice generously provided by Jerrold Turner (Harvard BWH)^34^. TcrdGDL mice^33^ were provided by Immo Prinz and Inga Sandrock (Hannover). GzmA/GzmB double knockout mice were provided by Todd Fehniger (WashU), and bones from GzmK and GzmB KO mice were obtained from David Granville (UBC). All studies were conducted in an Association of the Assessment and Accreditation of Laboratory Animal Care–accredited facility according to protocols approved by Rutgers New Jersey Medical School Comparative Medicine Resources. To induce γδ T cell depletion in TcrdGDL mice, 15 ng of diphtheria toxin (List Biological) per gram body weight was administered i.p. 24 and 48 hours prior to the initiation of the experiment.

### IEL isolation and flow cytometry

Small intestinal IELs were isolated as previously described^27^ and stained using fixable viability dye (eFluor 450 or 780, eBioscience), anti-CD3 (2C11, BioLegend), anti-TCRγδ (GL3, BioLegend), and anti-TCRβ (H57-597, BioLegend). Flow cytometry was performed on an LSR II (BD Biosciences) in the New Jersey Medical School Flow Cytometry and Immunology Core Laboratory and the data were analyzed by FlowJo (v. 10.4.2; Tree Star).

### Shedding assays

Mice were injected intraperitoneally with 10 mg/kg body weight of LPS from *E*. *coli* O111:B4 (Sigma-Aldrich) or 7.5 μg of recombinant TNF (Peprotech) and sacrificed after 90 minutes. Where indicated, mice were pre-treated with one of the following antibodies: 150 μg of anti-CD103 (M290, BioXcell) or rat IgG2a (RTK2758, BioLegend) 1h prior to LPS treatment; 200 μg of anti-NKG2D (HMG2D, BioXcell) or Armenian hamster IgG (BioXcell) 1h prior to LPS treatment; or 2 x 200 μg anti-TCRγδ (UC7-13D5, BioXcell) or Armenian hamster IgG (BioXcell) 24 and 48h prior to LPS treatment. To quantify shedding events, sections of formalin-fixed, paraffin-embedded small intestine from these mice were stained for cleaved caspase-3 (CC3) using immunohistochemistry (described below). Positive shedding events were identified via positive CC3 staining and cellular morphology (funnel formation and/or nuclear displacement).

### Intravital microscopy

Intravital imaging was performed as previously described^61,72^. Briefly, a loop of jejunum was exposed in anesthetized mice which had been administered i.v. Hoechst 33342. The anti-mesenteric surface of the intestinal mucosa was exposed and placed against a coverslip-bottomed 35 mm petri dish containing 1 μM Alexa Fluor 633. Time-lapse videos were captured using an inverted DMi8 microscope (Leica) equipped with a Yokogawa CSU-W1 spinning disk (Andor), a ×63/ 1.3 numerical aperture HC Plan APO glycerol immersion objective and an iXon Life 888 EMCCD camera (Andor). A DPSS 488 laser was used to image EGFP, DPSS 561 laser for DsRed, 640 nm diode laser for Alexa Fluor 633, and 405 nm diode laser for Hoechst dye. 15 μm z-stacks were acquired with 1.5 μm spacing with 20-30s between acquisition of a given z-plane. Imaris (v. 9.6, Bitplane) was used to render 3D reconstructions and apply an autoregressive tracking algorithm to enable quantification of IEL motility. Tracks were manually verified, and the track mean speed of each IEL was obtained. IEL contact with shedding cells was manually quantified and verified using both 3D maximum intensity projections and single slices of an entire z-stack. Dwell time indicates the duration of a single IEL/epithelial contact.

### Electron microscopy

Whole pieces of mouse jejunum were harvested and fixed overnight at 4°C in a solution of 2.5% glutaraldehyde, 2% paraformaldehyde, and 0.1M sodium cacodylate pH 7.4. Tissue was then post-fixed with 1% osmium tetroxide in 0.1 M sodium cacodylate buffer, stained en-bloc with 1% aqueous uranyl acetate, and dehydrated through a graded series of ethanol and propylene oxide. Next, tissues were incubated in a 1:1 mixture of EMBed 812 (Electron Microscopy Sciences 14120) and propylene oxide, then left in 100% EMBed 812 overnight before being embedded. Ultrathin sections (∼70 nm) were cut, and grids were stained with uranyl acetate and lead citrate. Sections were imaged using a Thermofisher Tecnai 12 transmission electron microscope and micrographs were recorded using a Gatan OneView 16-megapixel camera.

### Immunofluorescent and immunohistochemical staining

To visualize anti-CD103 antibody binding, 5 μm frozen sections of fixed mouse intestine were obtained as previously described^27^ and immunostained with Alexa Fluor 594-conjugated anti-Rat IgG to detect anti-CD103 antibody, along with Alexa Fluor 647-conjugated phalloidin and Hoechst 33342 (Invitrogen). Slides were mounted with Prolong Gold (Invitrogen).

To visualize granzymes and vesicles within IELs, small intestinal IELs were isolated, sort-purified, embedded on an 8 well chamberglass (Nunc) in 50% Matrigel (Corning), and incubated for 1h in RPMI-T medium^61^ containing 100U/ml interleukin (IL)-2 and 10ng/ml IL-15. IEL-containing Matrigel was fixed in 4% PFA, permeabilized in 0.5% Triton X100, and stained with primary antibodies against Granzyme A (BioLegend, PE conjugated), Granzyme B (BioLegend), LAMP1 (BioLegend). The appropriate secondary antibodies and Hoescht 33342 (Invitrogen) were applied, then samples were mounted in Prolong Gold (Invitrogen) and covered by a layer of mineral oil.

For all other immunostaining, intestinal tissue was fixed in 10% neutral buffered formalin, washed with 70% ethanol, paraffin-embedded and sectioned. 5 μm sections were deparaffinized, rehydrated, and blocked for endogenous peroxidases in 0.3% hydrogen peroxide, when applicable. Tissues were then subjected to antigen retrieval in either Tris- or citric acid-based antigen unmasking solutions (Vector Laboratories), blocked in 10% normal goat serum, and incubated with primary antibodies against CC3 (Cell Signaling Technology), ZO-1 (Santa Cruz) and/or GFP (Abcam). For immunofluorescent staining, fluorescent-conjugated secondary antibodies and Hoescht 33342 (Invitrogen) were applied, and slides were mounted in Prolong Gold. For immunohistochemical staining, sections were incubated with a horseradish peroxidase-conjugated anti-rabbit IgG secondary (Vector Laboratories), developed using a 3,3’-diaminobenzidine solution (Vector Laboratories), counterstained with hematoxylin, and mounted in Cytoseal XYL (Thermo Scientific).

Fluorescent images were captured on an inverted DMi8 microscope (Leica) equipped with a CSU-W1 spinning disk, ZYLA SL150 sCMOS camera (Andor), ×20/0.40 CORR, PL APO ×40/0.85 dry objectives, a x100/1.40-0.70 oil objective, and iQ3 acquisition software (Andor). Light microscopy was performed using a Keyence BZ-X710 using a x20/0.75 PL APO dry objective, and BZ-X Viewer acquisition software.

### ELISA

To measure intestinal TNF expression, whole pieces of mouse jejunum were homogenized with a bead beater in Bio-Plex Cell Lysis buffer (Bio-Rad). Protein concentrations were determined using the DC Protein Assay (Bio-Rad), lysates were diluted to a uniform protein concentration, and TNF levels were quantified using a mouse TNF ELISA kit (BioLegend) according to manufacturer instructions.

To measure granzyme secretion, sort-purified γδ or αβ IELs were seeded at approximately 400K cells per well in round-bottom 96-well plates coated overnight with 2 μg/ml anti-CD3 (145-2C11, BioLegend), 2 μg/ml polyclonal Armenian hamster IgG (BioXCell), and/or 2.5 μg/ml E-cadherin-Fc (Sigma). IELs were plated in RPMI-T medium containing 2 μg/ml PerCP-e710 conjugated anti-CD107a (1D4B, Invitrogen) for 90 min. Granzymes A and B secreted into the cell media were measured by ELISA (Abcam, Invitrogen). IELs were analyzed for CD107a externalization through flow cytometry, as described above.

### Quantitative PCR

Intestinal epithelial cells were isolated by inverting a segment of mouse jejunum on a feeding needle and incubated in Cell Recovery Solution (Corning). Epithelial cells were pelleted and resuspended in Trizol (Invitrogen) and homogenized using a 23½ gauge needle. RNA was isolated with a RNeasy kit (Qiagen) and cDNA was generated using the iScript cDNA Synthesis Kit (Bio-Rad). qPCR was performed on the QuantStudio 6 Real-Time PCR System (Applied Biosystems) using PowerUp SYBR Green Master Mix (Applied Biosystems) and primers for *H60* (forward, 5’-GAGCCACCAGCAAGAGCAA-3’; reverse, 5’-CCAGTATGGTCCCCAGATAGCT-3’), *Rae1* (forward, 5’-ATCAACTTCCCCGCTTCCA-3’; reverse, 5’-AGATATGAAGATGAGTCCCACAGAGATA-3’), *Ulbp1* (forward, 5’-TTCACATAGTGCAGGAGACTAACACA-3’; reverse, 5’-ACTGGCCACACACCTCAGC-3’) (Li et al. 2012), or *Gapdh* (forward, 5’-AAGGTGGTGAAGCAGGCATCTGAG-3’; reverse, 5’-GGAAGAGTGGGAGTTGCTGTTGAAGTC3’). Fold changes were calculated using Ct values normalized to *Gapdh* expression.

### Generation of bone marrow chimeras

8-week-old Tcrd KO mice were exposed to 11 Gy of gamma irradiation. 24 hours later, irradiated mice were retro-orbitally injected with 5×10^6^ bone marrow cells from donor mice. Tissue was harvested 8 weeks post-engraftment.

### Statistical analyses

Statistical analysis was performed using GraphPad Prism software. Two-tailed unpaired Student *t* tests were used to directly compare two independent samples; a p≤ 0.05 was considered statistically significant. Comparisons between multiple independent variables were performed using one-way ANOVA followed by post-hoc Tukey multiple comparisons tests for pairwise comparisons. The two-stage Benjamini, Krieger, and Yekutieli procedure was used post hoc to control the false discovery rate. All data are presented as either the mean ± SEM or with a 95% confidence interval.

## Abbreviations

BM: basement membrane
CC3: cleaved caspase-3
DT: diphtheria toxin
Gzm: granzyme
IBD: inflammatory bowel disease
IL: interleukin
IEL: intraepithelial lymphocyte
KO: knockout
LIS: lateral intercellular space
LPS: lipopolysaccharide
TNF: tumor necrosis factor
TCR: T cell receptor
ZO-1: zona occludens-1

## Acknowledgments

The authors would like to thank Jerry Turner for providing the mRFP-ZO-1 Tg mice, Juan Flores for assistance with embedding and Raj Patel for thin sectioning for electron microscopy. We also gratefully acknowledge Angela Patterson for her thoughtful input. Electron microscopy was performed in the AICF at Rutgers-Newark. Cell sorting was performed at the NJMS Flow Cytometry and Immunology Core Laboratory and supported by National Institute for Research Resources Grant S10RR027022. This work was supported by National Institute of Health Grants R21AI143892 (K.L.E), T32AI125185, F30DK121391 (M.D.H.) and the New Jersey Health Foundation (K.L.E.) and grants BB/J004529/1 and BB/K018256/1 (A.J.M.W.). D.J.G. was funded by a Canadian Institutes for Health Research Foundation Grant.

## Author contributions

M.D.H. designed and performed experiments and wrote the manuscript. N.B.G. designed and performed experiments and reviewed the manuscript. T.J.K, J.A., and E.M.B. performed experiments. M.R.Z., I.S., I.P. and D.J.G. provided resources for the study and reviewed the manuscript. A.J.M.W. conceived the study, contributed to experimental design and reviewed the manuscript. K.L.E. conceived the study, performed experiments, supervised the research and wrote the manuscript.

## Video Legends

**Supplemental Video 1**. Intravital imaging of γδ T cells (green), luminal Alexa Fluor 633 (red) and nuclei (white) in jejunum of a TcrdEGFP mouse treated with 10mg/kg LPS for 90 min. Frames were collected approximately 30 s.

**Supplemental Video 2**. Intravital imaging and 3D reconstruction of a γδ T cell (green) directly interacting with a ZO-1 funnel (white) in the jejunum of a LPS-treated TcrdGDL; mRFP-ZO-1 Tg mouse. Frames were collected approximately 17 s.

**Figure S1.**
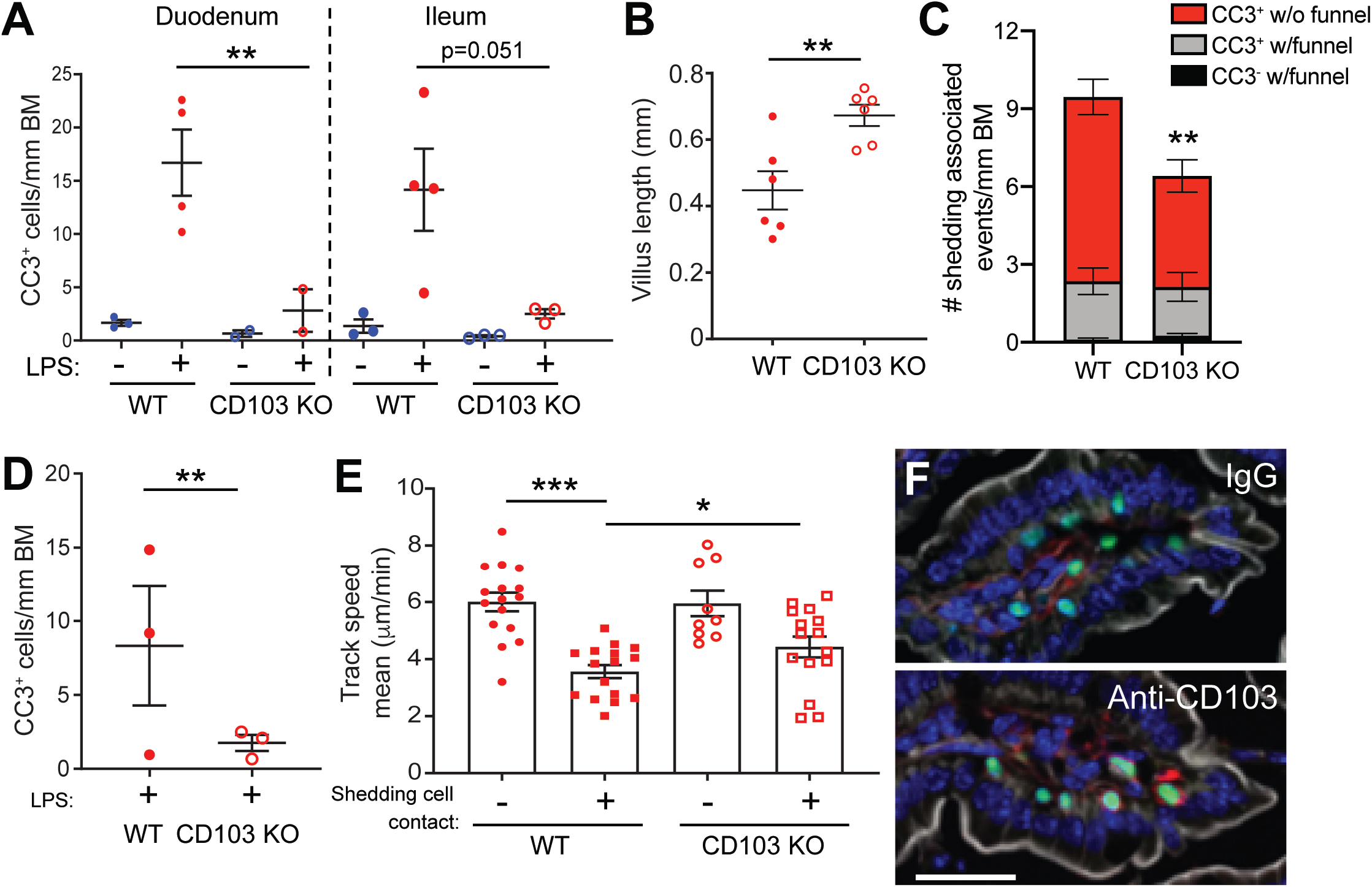
γδ CD103 regulates γδ IEL migratory behavior to facilitate LPS-induced cell shedding independent of the microbiome. (A) Quantification of CC3^+^ shedding events in the duodenum and ileum of LPS-treated WT and CD103 KO mice. (B) Villus length in WT and CD103 KO mice. (C) Number of events observed during various stages of cell shedding in LPS-treated WT and CD103 KO mice. n = 8-10 mice. (D) Number of CC3+ shedding events in the jejunum of cohoused WT and CD103 KO mice following LPS exposure. (E) Mean track speed of WT or CD103 KO γδ IELs that do or do not contact shedding cells (9-17 tracks). (F) Immunostaining for rat IgG (red), showing anti-CD103 on the surface of γδ IELs (green). White: F-actin. Scale bar = 25 μm. N = 3-6 mice; *p<0.05, **p<0.01, ***p<0.001.

**Figure S2.**
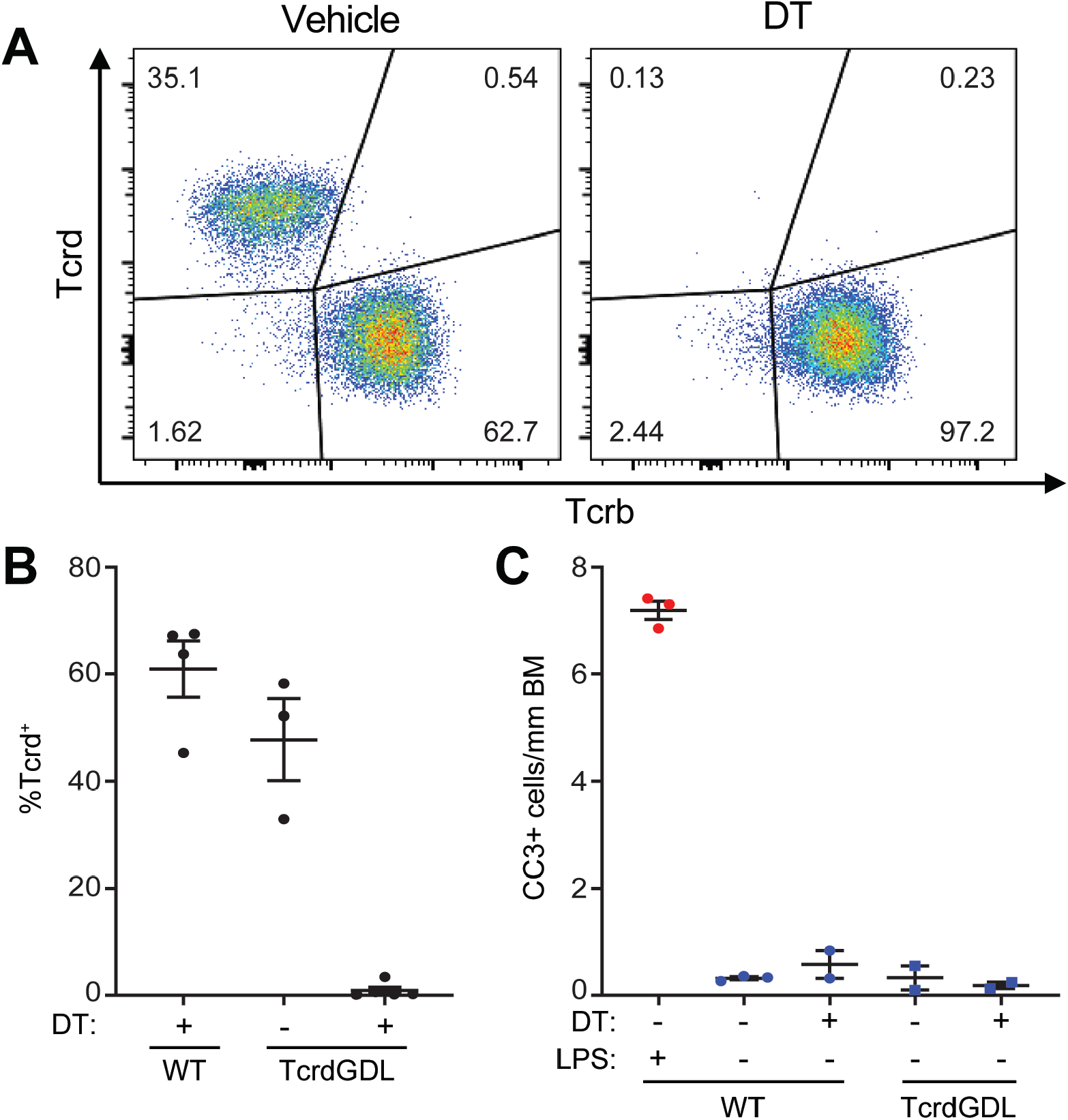
Diphtheria toxin effectively depletes γδ IELs in TcrdGDL mice. WT or TcrdGDL mice were treated with diphtheria toxin (DT) or vehicle control prior to PBS or LPS treatment. (A) Representative flow cytometry plots showing the (B) percent of TCR γδ^+^ IELs in the small intestine, gated on live CD3^+^ cells. (C) Quantification of CC3+ shedding events in jejunal tissue, n = 2-5 mice.

**Figure S3.**
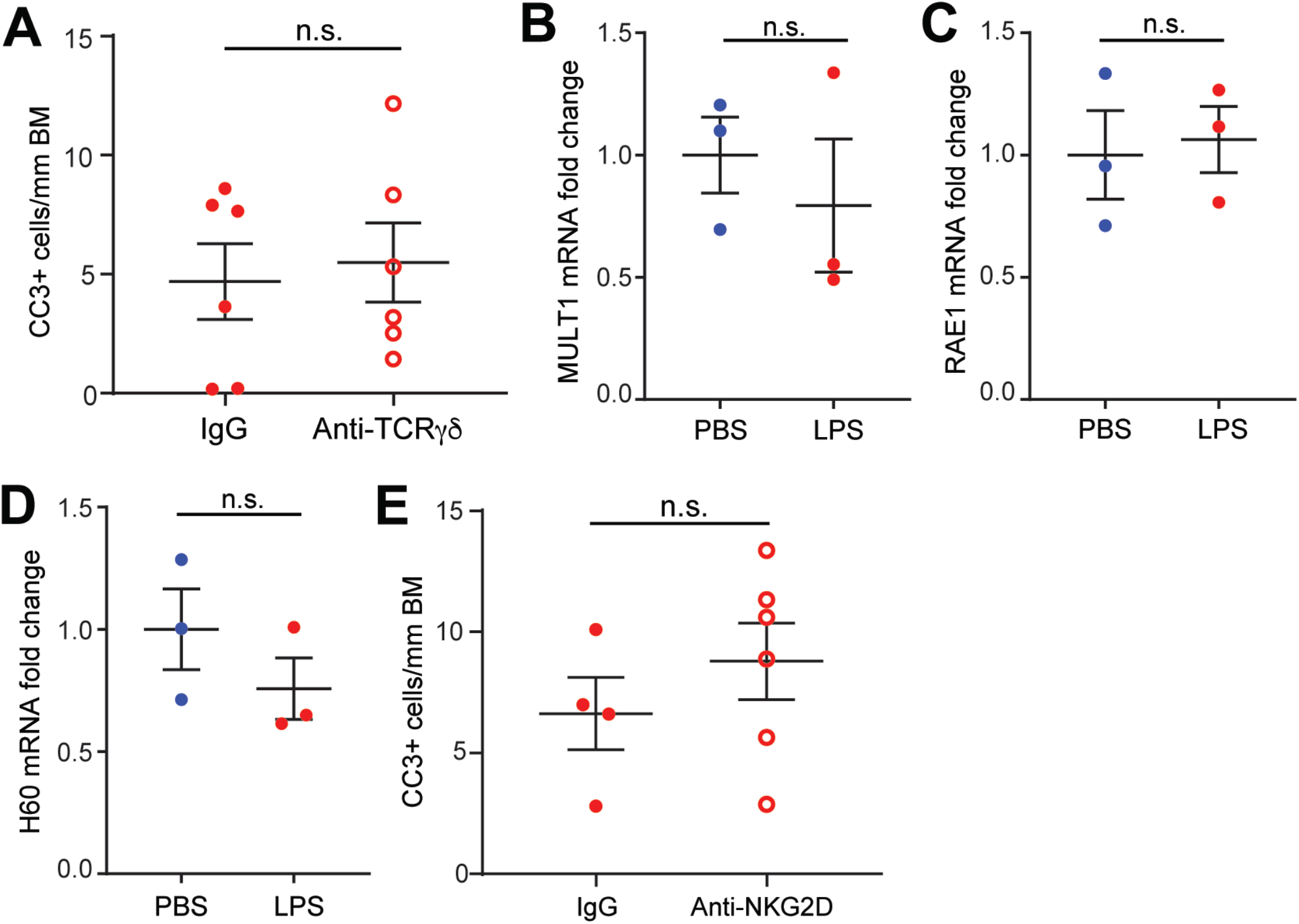
Signaling through TCRγδ or NKG2D is not required for LPS-induced cell shedding. (A) Quantification of CC3 staining in mice treated with anti-TCRγδ or IgG control prior to LPS administration. (B-D) RNA expression of NKG2D ligands (MULTI, RAE1, and H60) in jejunal epithelium isolated from PBS- or LPS-treated WT mice. Data are shown as the fold change in expression relative to control mice. (E) Quantification of CC3 staining of jejunum from mice administered NKG2D blocking antibody or IgG control prior to LPS exposure, n = 3-6 mice; n.s., not significant.

**Figure S4.**
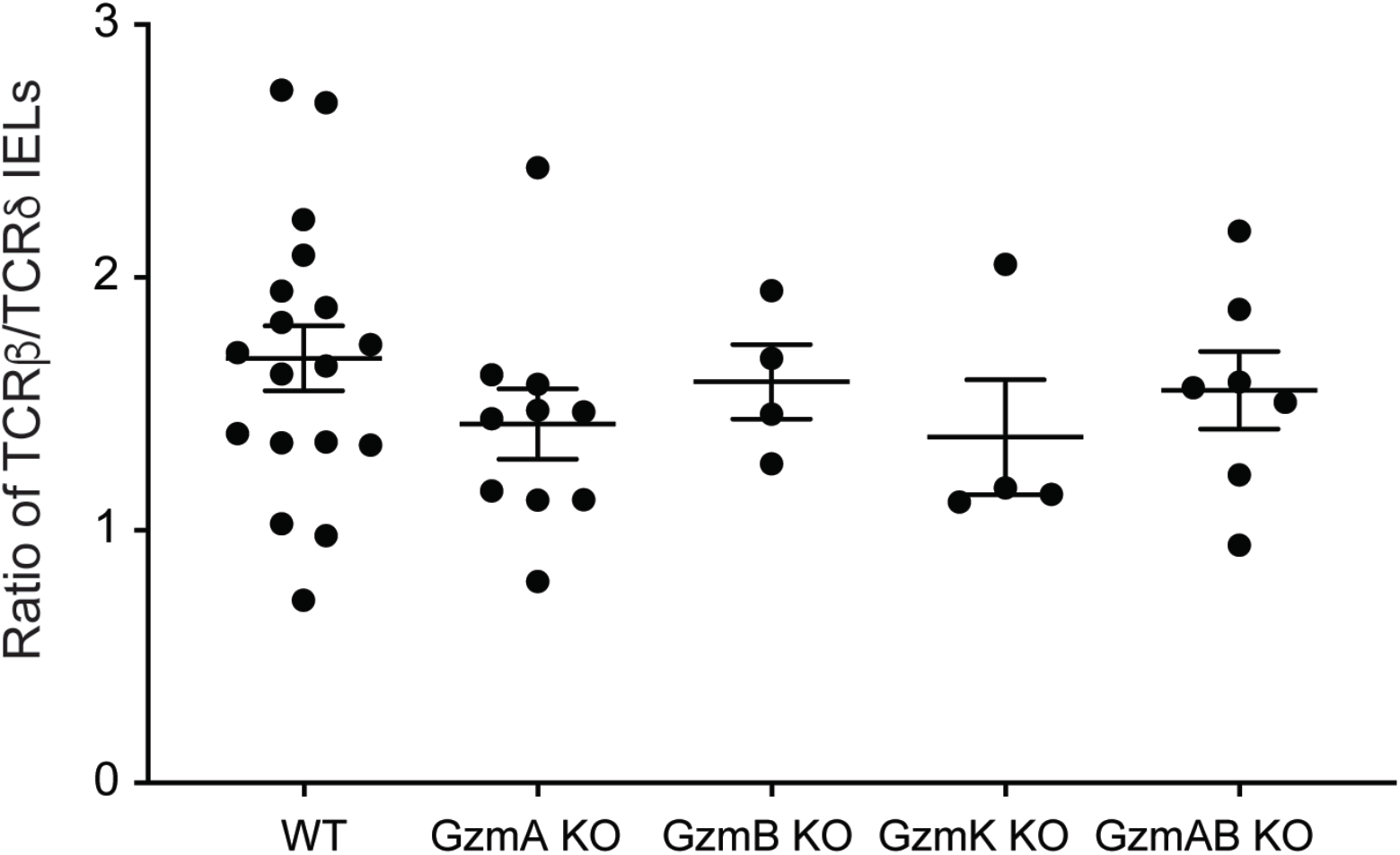
IEL populations are comparable between wildtype and granzyme-deficient bone marrow chimeras. Ratio of TCRβ^+^ to TCRδ^+^ IELs in whole small intestine as determined by flow cytometry. Gated on live CD3^+^ cells. n = 4-15 mice.

